# In-depth analysis of *Klebsiella aerogenes* resistome, virulome and mobilome worldwide

**DOI:** 10.1101/2023.08.15.553288

**Authors:** Sergio Morgado, Érica Fonseca, Fernanda Freitas, Raquel Caldart, Ana Carolina Vicente

**Affiliations:** Laboratório de Genética Molecular de Microrganismos, Instituto Oswaldo Cruz, Rio de Janeiro 21040-360, RJ, Brazil; Centro de Ciências da Saúde, Universidade Federal de Roraima, Boa Vista 69300-000, RR, Brazil

## Abstract

*Klebsiella aerogenes* is an emergent pathogen associated with outbreaks of carbapenem-resistant strains. To date, studies focusing on *K. aerogenes* have been small-scale and/or geographically restricted. Here, we analyzed the epidemiology, resistome, virulome, and mobilome of this species based on 561 genomes, spanning all continents. Furthermore, we sequenced four new strains from Brazil (mostly from the Amazon region). Dozens of STs occur worldwide, but the pandemic clones ST93 and ST4 have prevailed in several countries. Almost all genomes were clinical, however, most of them did not carry ESBL or carbapenemases, instead, they carried chromosomal alterations (*omp*36, *amp*D, *amp*G, *amp*R) associated with resistance to β-lactams. Integrons were also identified, presenting gene cassettes not yet reported in this species (*bla*IMP, *bla*VIM, *bla*GES). Considering the virulence loci, the yersiniabactin and colibactin operons were found in the ICEKp10 element, which is disseminated in genomes of several STs, as well as an incomplete salmochelin cluster. In contrast, the aerobactin hypervirulence trait was observed only in one ST432 genome. Plasmids were common, mainly from the ColRNAI replicon, with some carrying resistance genes (*mcr, bla*TEM, *bla*NDM, *bla*IMP, *bla*KPC, *bla*VIM) and virulence genes (EAST1, *sen*B). Interestingly, plasmids containing the colicin gene occurred in hundreds of genomes of different STs.

## Introduction

The bacteria formerly known as *Enterobacter aerogenes* has recently been renamed *Klebsiella aerogenes* based on whole-genome sequence phylogenetics [1,2]. This species is a ubiquitous member of the Enterobacteriaceae family, and although it has always been considered an opportunistic pathogen, recently, interest in this organism has increased, mainly due to the emergence of multidrug-resistant and carbapenem-resistant strains [3]. This may be due to the acquisition of antibiotic resistance genes carried by mobile elements, such as plasmids and conjugative integrative elements, which have already been identified in *K. aerogenes* [4-6). However, in this species, the main mechanisms of resistance to carbapenems are attributed to chromosomal overexpression of AmpC β-lactamase and mutations that affect membrane permeability [4]. Furthermore, reports show that polymyxin resistance is occurring in strains of this species due to mutations in some housekeeping genes (*mgr*B, *pmr*A, *pmr*B, and *ept*A) [7,8] and, eventually, by plasmids carrying the *mcr* gene.

There are continuous reports of the high frequency of outbreaks of this emergent pathogen, some of them associated with high mortality rates [9, 10], in different clinical settings around the world, such as neonatal [11-13], geriatric [14], and intensive care units [8, 15]. Thus, *K. aerogenes* was included in the ESKAPE group, which encompasses important pathogens associated with antimicrobial resistance [5]. Among the hundreds of known *K. aerogenes* lineages, comparative genomics showed that type 4 (ST4) and ST93 lineages likely represent the dominant lineages associated with human infections worldwide [4]. In addition, some lineages have been associated with virulence determinants, such as siderophores and toxins, which potentiate their persistence and favor the emergence of nosocomial outbreaks [4, 5, 8, 12].

So far, studies focusing on *K. aerogenes* have been small-scale and/or with a restricted geographic perspective [4, 5, 16]. Here, in addition to generating new genomes of this species in Brazil and contextualizing them in the global scenario, we performed an in-depth analysis of the resistome, virulome, and mobilome of this emergent pathogen.

## Results

### *K. aerogenes* sequence typing and epidemiology

In total, 561 *K. aerogenes* were analyzed, where most (n=477) were associated with human infections, and few were associated with the environment (n=15) and animals (n=13), covering the period from 1955 to 2022. For all sources, several STs were identified, where ST93 (n=121) and ST4 (n=57) were the most prevalent (Figure 1, Table S1). Other STs were present in no more than 20 genomes, showing that the prevalence of ST93 and ST4 is indeed much higher (Table S1). The phylogeny showed the division of the genomes into four main clusters, with the main one encompassing most of the genomes (493/561) into several subclusters, including ST4 and ST93 (Figure 1). Dozens of genomes (n=41) did not have any ST assigned, showing the genetic diversity of this species. Therefore, we submitted these allelic patterns to pubMLST database (https://pubmlst.org/bigsdb?db=pubmlst_kaerogenes_seqdef), where 41 new ST profiles were generated and included in this study (ST398-ST437, and ST440). The ST4 genomes are from 2004 to 2022 and occur in several countries in South America, North America, Asia, and, only in the United Kingdom in Europe. Most of this ST is associated with human sources; however, one genome is from the environment (Table S1). ST93 is more widespread than ST4 genomes as it also occurs in several European countries and Central America. Most ST93 genomes are from humans, occurring from 1997 to 2022 (Table S1).

**Figure 1.**
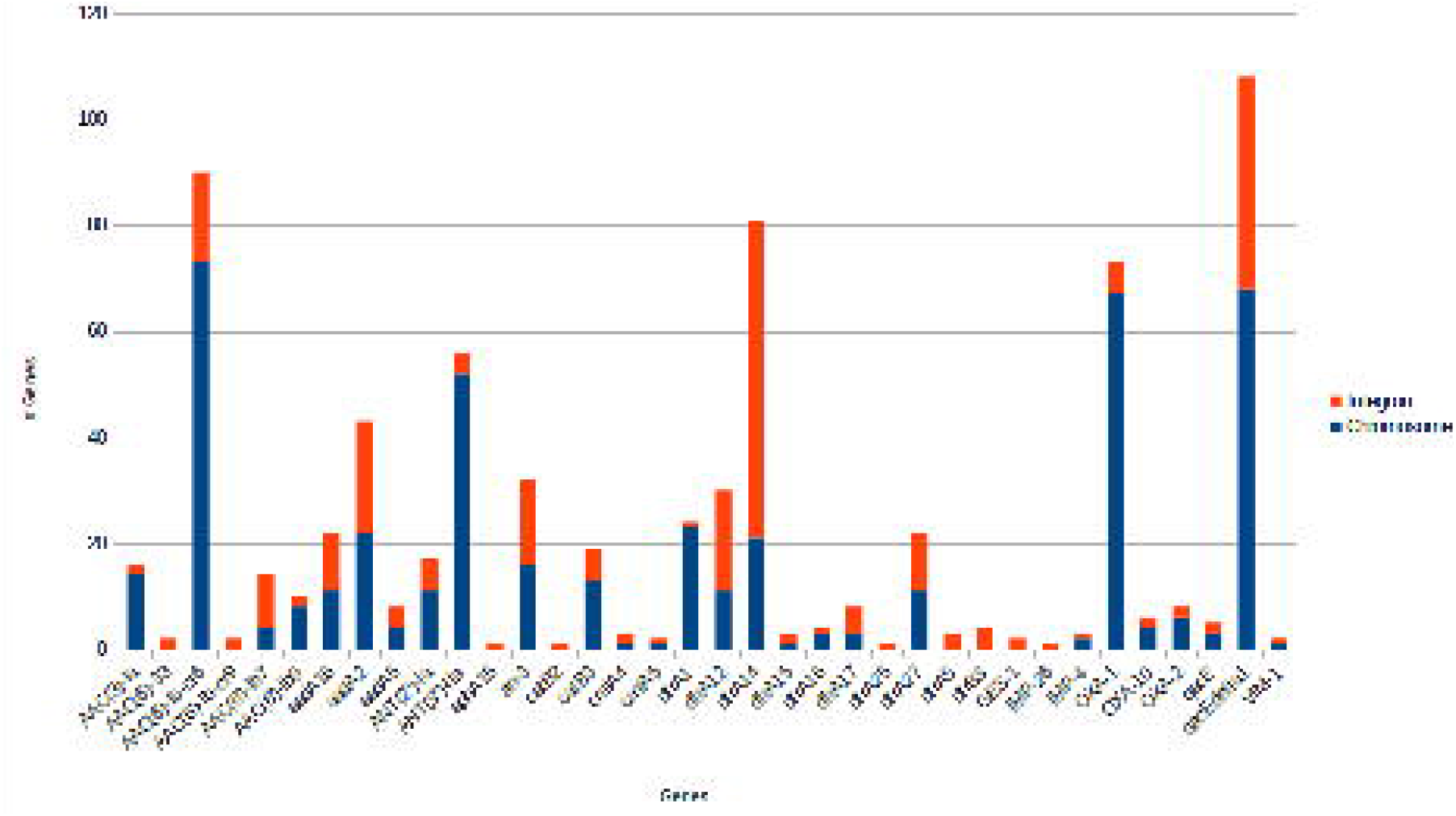
Maximum-likelihood tree based on the core genome of *K. aerogenes*. The ST number of each genome is next to the accession number. There are two orbits of colored blocks, where the innermost represents the source of isolation, and the outermost represents the regions of the genomes. Next, five colored circles indicate genomes with yersiniabactin (green), colibactin (beige), integron (purple), cloacin (blue), mutations in AmpD, AmpR, Omp35/36 (red). More external to these circles, the different sizes of the bars (0 to 5) indicate the antibiotic resistance (red) and virulence (blue) scores of the genomes. The genomes sequenced here are in green background color. Red circles at branches indicate bootstrap values >70%.

In Brazil, *K. aerogenes* genomes are associated with human infections from 2006 to 2016 in three states in the Southeast region (Minas Gerais, Paraná and São Paulo), where three STs were observed: ST93 (11/20), ST16 (2 /20) and ST4 (3/20) (Table S1). Furthermore, the new genomes sequenced here came from other states in the Southeast region, Rio de Janeiro (ST296), and from the North region, Roraima (ST93 and ST440). ST296 and ST440 represent two new STs in Brazil.

### Brazilian *K. aerogenes* antibiotic susceptibility

The four Brazilian strains of *K. aerogenes* were tested for various antibiotics, including carbapenems and cephalosporins (Table 1). The Ka-04RR strain showed resistance to a greater number of antibiotics, mainly cephalosporins and β-lactams. The punctual pattern of resistance to carbapenems, cephalosporins and β-lactams suggests the absence of an enzymatic resistance mechanism. Even so, all strains would be considered multidrug resistant (MDR) [17].

**Table 1.**
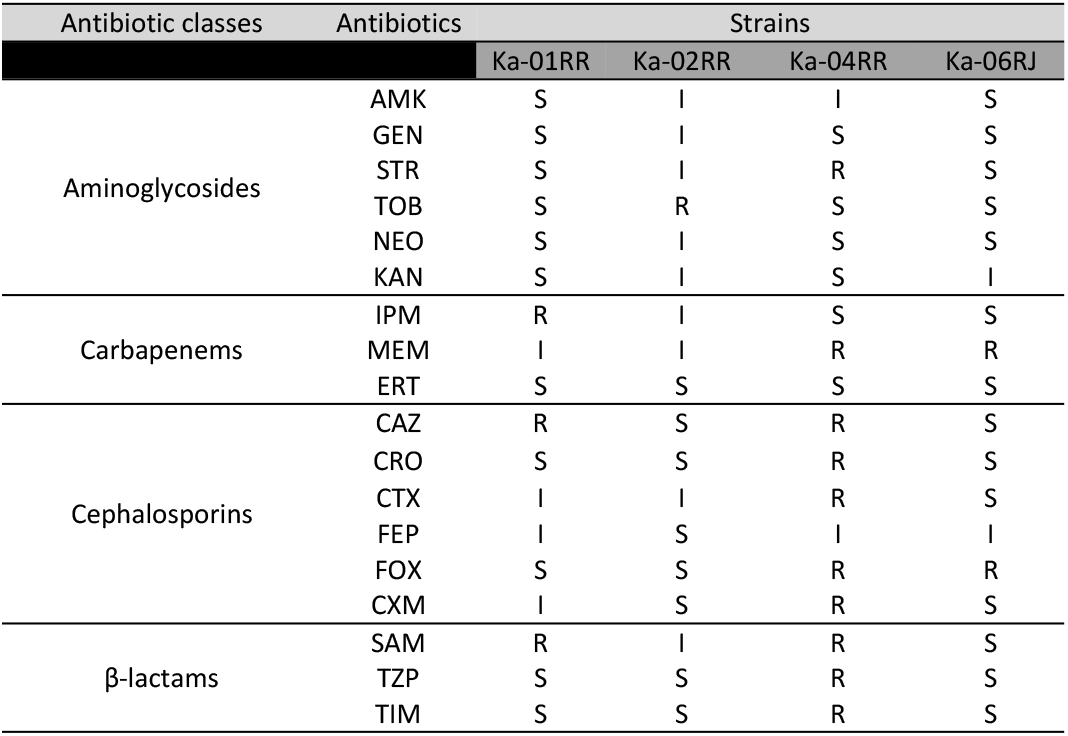
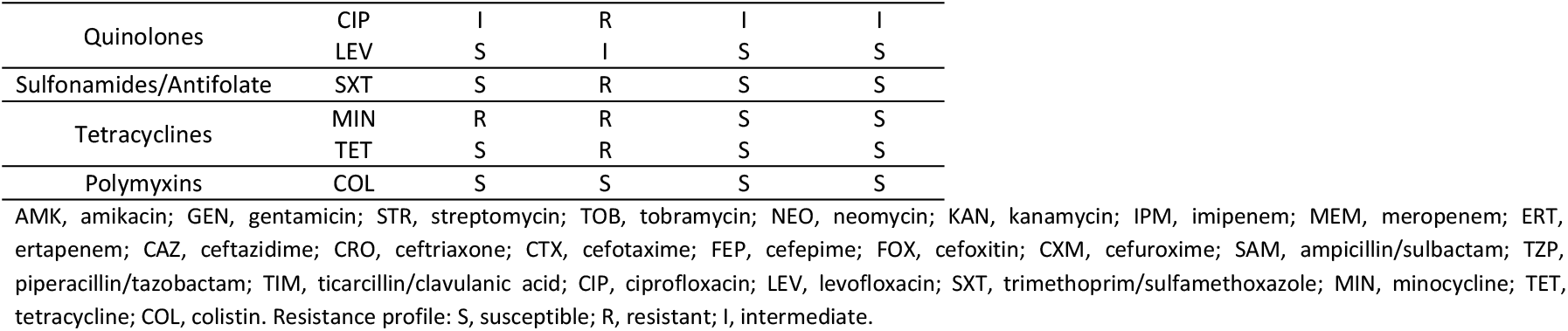
Antibiotic susceptibility of Brazilian *K. aerogenes*.

### *K. aerogenes* resistome analysis

Based on Kleborate’s metrics for antibiotic resistance, 366 genomes had score 0; 42, score 1 (ESBL producers); 150, score 2 (carbapenemase producers); and 3, score 3 (carbapenemase producers plus colistin resistance). The most prevalent carbapenemases were *bla*KPC (−2 and -3), *bla*OXA-48 and *bla*NDM (−1, -5, -7 and -9) (Table S1). Therefore, although most genomes are of clinical origin, most of them do not carry carbapenemases. Hundreds of genomes (n=169; 30%) carried genes associated with three or more classes of antibiotics, characterizing them as MDR, where most of these belonged to ST93 (n=43), ST4 (n=14) and ST116 (n=11) (Table S1). In contrast, there are also several genomes of pandemic clones ST4 (n=39) and ST93 (n=71) that lack acquired resistance genes (Table S1). Likewise, the Brazilian genomes sequenced here also did not show acquired resistance genes, including the ST93 (Ka-04RR), although this strain is MDR.

Only 12 genomes were predicted to harbor the *mcr* gene, where nine were related to ST56 in China (Table S1). Moreover, analyzing PmrA, three Dutch ST364 genomes were observed carrying the G53S amino acid substitution (Table S2), which has been associated with colistin resistance [7]. The GCA_000950145 genome showed a large block of dozens of amino acid replacements from position 85 to 109, including insertions and deletions (Table S2), which could also affect its role. Interestingly, the L162M amino acid replacement was observed in almost all genomes (551/561), even in some colistin-susceptible strains (Table S2). Furthermore, three genomes had nonsense mutations in MgrB (GCA_021837555, GCA_021837885, GCA_022016415), also being associated with colistin resistance [18].

*In vitro* analyses showed that Ka-04RR was resistant to meropenem and to most cephalosporins and β-lactams tested (Table 1). However, this strain lacked acquired ESBL and carbapenemases genes (Table S1). Thus, we explored amino acid replacements at chromosomal loci associated with β-lactam resistance in this strain, in addition to the entire genomic dataset. We focused on AmpC and outer membrane porins (Omp35 and Omp36), including their associated regulatory proteins (AmpD, AmpG, AmpR, and OmpR). Comparisons were evaluated using wild-type sequences from carbapenem-susceptible strains. All respective genes (*amp*C, *omp*35, *omp*36, *amp*D, *amp*G, *amp*R, and *omp*R) were identified in most genomes (Table S3). They showed different patterns of single nucleotide polymorphisms (SNPs), some leading to premature translation stops in their respective proteins. The AmpD protein was truncated in 44 genomes, including Ka-04RR. Mutation analysis, considering all genomes, showed that 65% (120/187 sites) of amino acid sites were conserved and several replacements were common, such as V134A, V135A, Q138R (Table S4). Furthermore, replacements associated with carbapenem resistance (P39S, A94T, W95L, S112L, I113S, I160V, R161H [4]) were observed in several genomes, most of them belonging to ST4 and ST93 (Table S4). The AmpG sequences showed 91% (451/491 sites) of conserved amino acid sites (Table S5), where four genomes had truncated proteins, including the Ka01-RR genome, while the Ka02-RR genome lacked *amp*G. Also, several mutational patterns were observed, such as L89I, I93V, T424I, L430V, G461A and I479V (Table S5). Even so, all known AmpG activation motif residues (G25, A122, Q124, A181 [19]) were conserved, except for GCA_021937655 (A181V). AmpR was found truncated in two genomes and absent in four genomes (GCA_000534135, GCA_000950145, GCA_011604725, GCA_022759585), which also lacked *amp*R. Overall, the genomes showed 87% (255/292 sites) of conserved amino acid sites (Table S6). AmpR sites associated with its role (R86, G102, D135 [20, 21] were also conserved except for D135A/N substitutions in four genomes (GCA_900558125, GCA_025115605, GCA_022016415, GCA_021936135) (Table S6). Regarding the porins, Omp35 was truncated in 21 genomes, but well conserved in the remaining genomes (93% of amino acid sites conserved; 337/359 sites) (Table S7). Most ST202, ST296, ST300, ST375 genomes had common mutations: S182V, D225N, D259N, H260Y (Table S7). Omp36 was the analyzed protein with the highest rates of nonsense mutations, where 98 genomes had premature stop codons. Among the Omp36 positive genomes, several sites had mutations, which resulted in 72% (270/375 sites) of conserved sites (Table S8). Most mutations were concentrated in the final half of the protein (Table S8). Some replacements were frequently found in ST-specific genomes, such as I59V in ST4, ST231, ST237; D189G in ST4, ST16, ST93; D205E in ST93 and ST228 (Table S8). The OmpR showed the highest conservation among the proteins explored with 97% (233/239 sites) of conserved amino acid sites (Table S9). Two important sites (G63 and R150 [22, 23]) were conserved in all genomes.

Among the proteins analyzed, amino acid replacements and/or premature translation stops in AmpD, AmpR, Omp35, and Omp36, which can lead to a phenotype of greater resistance to β-lactam, were present in 148 genomes spread across the phylogeny, regardless the ST (Figure 1). Most of these genomes (117/148; ∼80%) did not contain ESBL (19/148) and/or carbapenemase (12/148) genes, suggesting that these mutations are selection mechanisms in the absence of enzymatic resistance genes. Even more so because most of these genomes (100/148; ∼67%) also lacked acquired resistance genes (Table S1). Thus, contrasting to the low prevalence of acquired genes, a large proportion of genomes carrying chromosomal alterations associated with resistance to β-lactam was identified. Therefore, without this deep analysis, the *in silico* inference of β-lactam resistance in *K. aerogenes* would be underestimated.

Hundreds of class 1 integrons (n=139) could be identified in 96 genomes, mainly from ST93 (n=29) (Table S10). Different combinations of gene cassettes were present, where most integrons presented the *dfr*A gene, while a smaller proportion presented *qac*E, *ant, aac, aph, arr, bla*OXA, and *bla*GES, some of them presenting several alleles. Furthermore, *bla*VIM and *bla*IMP carbapenemase genes were also present in gene cassettes (Table S10 and Figure 2).

**Figure 2.**
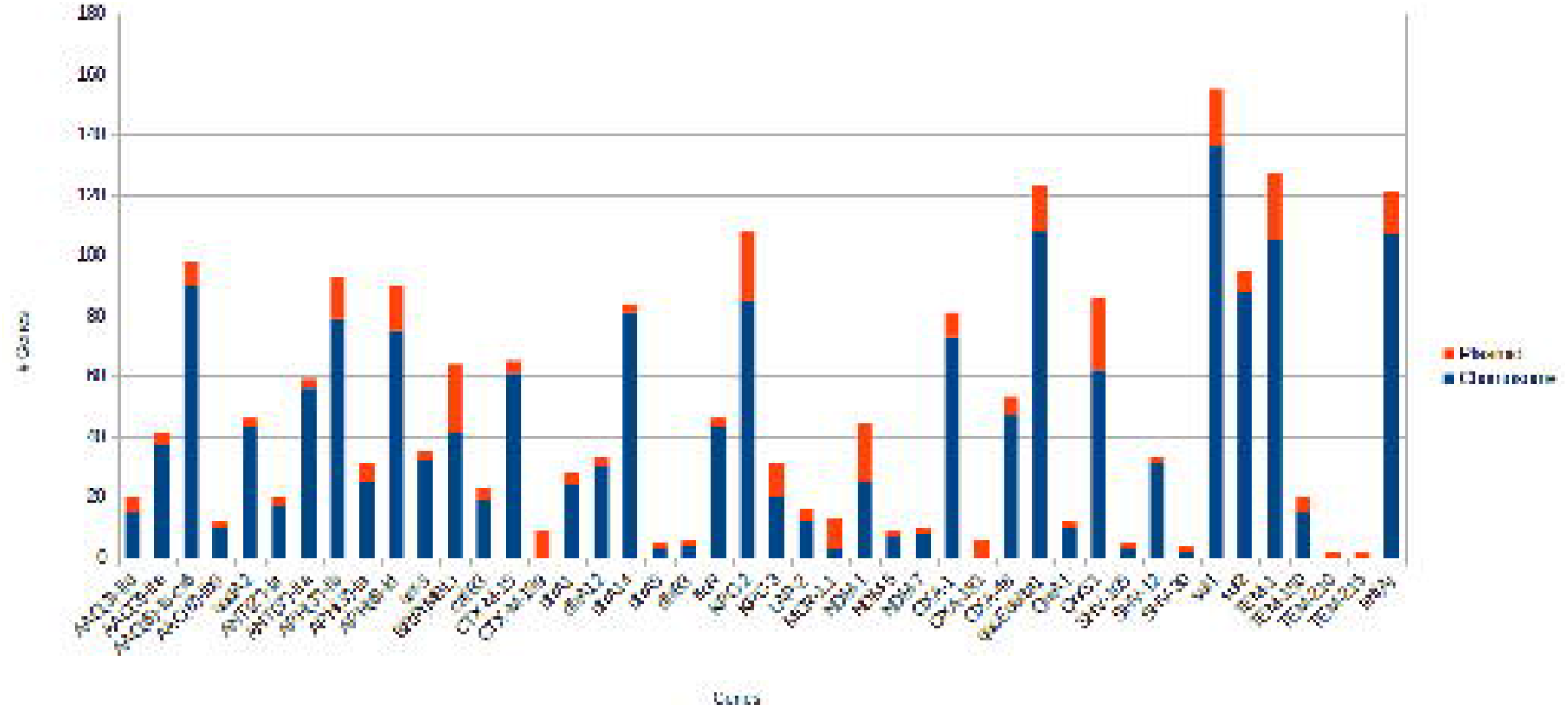
Number and proportion of antibiotic resistance genes in the context of integrons and chromosomes.

### *K. aerogenes* virulence analysis

Regarding the analysis of virulence loci, 322 genomes (57%) had a score of 0 (no *ybt* or *clb* loci); 16, score 1 (*ybt* only); 222, score 2 (*clb* positive); and 1, score 3 (aerobactin positive)(Table S1). All genomes with score 2 co-carried the *ybt*. Furthermore, among the major STs, these loci were prevalent in the genomes of pandemic clones ST93 (n=114) and ST4 (n=47) (Table 2). Almost all *ybt* and *clb* loci were in the context of ICEKp10 (208/238 *ybt*+ genomes), regardless of ST, with the majority of ST4 having the *ybt* type 17 (n=45), and majority of ST93 having the *ybt* type 17 (n=59) and *ybt* type 20 (n=52) (Table S1), while only the *clb* type 3 was found associated with these loci. ICEKp other than ICEKp10 were identified in a few genomes and carried only the *ybt*, lacking *clb* (Table S1). Almost all Brazilian genomes of *K. aerogenes* had a virulence score of 2, with the majority having *ybt* and *clb* loci in the ICEKp10 context. Only one environmental genome (ST93; score 2) presented the *ybt* 17 and *clb* 3 loci in the context of ICEKp10, while other animal and environmental genomes lacked these virulent loci. Curiously, the salmochelin (*iro*) virulence trait, which is not considered in the scoring system, is present in almost all *K. aerogenes* genomes (517/561; ∼92%) (Table S1). However, no genome presented the entire cluster (*iro*BCDEN), lacking *iro*DE. In addition, although lacking the *ybt* and *clb* loci, most animal and environmental genomes carried the *iro* loci (19/27; ∼70%). Another prevalent virulence factor among the genomes was the T6SS, present in 553/561 (∼98%). Regarding toxins, two factors (EAST1 and *sen*B) were found in the genomes of ST240, ST15, ST93, ST331, ST364. In short, ICEKp10 carrying the *ybt* and *clb* is widespread in dozens STs, prevailing in ST4 and ST93.

**Table 2.** Prevalence of virulence loci in the main STs of *K. aerogenes*.

| ST | # | *ybt* | *clb* | *iro* | T6SS |
| --- | --- | --- | --- | --- | --- |
| 93 | 121 | 114 (94%) | 114 (94%) | 121 (100%) | 118 (97%) |
| 4 | 57 | 47 (82%) | 47 (82%) | 57 (100%) | 55 (96%) |
| 135 | 16 | 0 | 0 | 16 (100%) | 16 (100%) |
| 116 | 13 | 1 (7%) | 1 (7%) | 13 (100%) | 13 (100%) |
| 197 | 12 | 8 (66%) | 8 (66%) | 12 (100%) | 12 (100%) |
| 56 | 10 | 0 | 0 | 10 (100%) | 10 (100%) |

In four genomes (GCA_020982565.1, GCA_001631645.1, GCA_003952125.1, GCA_021902315.1), the *ybt* type 4 was predicted, this type being associated with plasmid origin (Table S1). However, analyzing these genomes, we were unable to identify the plasmid replication genes. This could be due to genome assembly fragmentation, however, the *ybt* loci of these genomes were close to integrases, and flanked by tRNA-Asn (in 3/4 genomes), which is a feature associated with chromosomal insertion. Thus, if *ybt* type 4 is on a mobile element, it appears to have the ability to integrate into the chromosome.

The US clinical genome GCA_023553355.1 (ST432) was the only one to show a virulence score of 3 due to the presence of the aerobactin loci (*iuc* 1; *iuc*A-D and *iut*A) in addition to the hypermucoviscosity-associated gene *rmp*A2 (Table S1). These loci likely lie within a genomic island (contig ABGJKZ010000015.1), in a region of approximately 75 kb length (7374 bp – 82380 bp) flanked by repetitive regions (93 bp repeats with 98% identity) and three transposase genes (ISL3-like element, ISNCY-like element). Furthermore, this region has 99% identity and 90% coverage with several *Klebsiella pneumoniae* plasmids (e.g., pFQ61_ST383_NDM-5, pAP855, pVir-CR-hvKP-C789), some of them designated as hypervirulent plasmids. Curiously, although only this genome contained the *iuc*A-D loci, most genomes (n=552; 98%) harbored the *iut*A gene, which encodes the aerobactin receptor.

### *K. aerogenes* plasmid resistome/virulome

Plasmids were predicted (n=888) in 391 genomes (391/561; 69%), based on the presence of plasmid replicon genes (*rep*), being the ColRNAI replicon the most common. Plasmids ranged per genome from 1 to 10, with a median of 2 plasmids. The plasmids had a median size of 21.5 kb and 51% GC content (Table S11). Considering the mobility of the plasmids, 64 would be conjugative (48.7 kb median size, 49% GC content), as they carried relaxase, *vir*D4, and *vir*B4 homologues genes [24]; 450 would be mobilizable (10.2 kb median size, 53% GC content) due to the presence of oriT sequences and/or relaxase genes, without T4SS-like genes; and 374 would be non-mobilizable (19.9 kb median size, 49% GC content) (Table S11). Of note, considering the contents of these plasmids, the cloacin gene (a type of colicin) and its immunity gene were identified in hundreds (n=172; Figure 1) of small mobilizable plasmids (up to 19 kb, median size of 9.2 kb). These plasmids were mainly harbored in the ST93 (n=93) and ST4 (n=34) genomes, but were also present in dozens of other STs in smaller numbers (n<10). These genomes are from several continents, presenting a wide time scale (2002 to 2021).

Only 16/888 plasmids (in 13 genomes from different STs), with a median size of 98.6 kb, harbored virulence genes, most of these having only one gene, where *tra*J (associated with invasion), EAST1 (exotoxin), and *sen*B (exotoxin) genes were the most prevalent (Table S12). In addition, a putative plasmid (CDAGCNO010000015) carried four *fim* genes (*fim*C, *fim*D, *fim*F, *fim*G), but, curiously, it was predicted as non-mobilizable. None of the conjugative plasmids carried virulence genes.

Considering resistance genes, 155/888 plasmids (in 110 genomes), with a median size of 46.1 kb, carried 1-12 genes (median of two genes) (Table S13), associated to several classes of antibiotics, including cephalosporins (*bla*TEM, *bla*CTX-M, *bla*OXA), carbapenems (*bla*NDM, *bla*IMP, *bla*KPC, *bla*VIM), aminoglycosides (*aac, aad, ant, arm*A, *aph*A), phenicol (*cat*), sulfonamides (*sul*), macrolides (*mph*A, *msr*E), quinolones (*qnr*S). The most prevalent resistance genes in plasmids were *aph*(6)-Id (present in 15 plasmids); *bla*NDM-1 (19); *bla*TEM-1 (22); BRP (gene encoding bleomycin resistance protein) (23); *bla*KPC-2 (23); *qnr*S1 (24), where the majority were scattered in genomes of different STs (Table S13 and Figure 3). Importantly, the *mcr* gene (colistin resistance) was associated with plasmids (10/12 genomes with *mcr*). These genomes belonged to ST56/China (majority; IncI2 replicons) and ST207/Singapore (IncX4 replicons), in which eight co-harbored the CTX-M-199 gene (Table S13 and Figure 3). Moreover, most of these plasmids were predicted as conjugative (Table S13). Interestingly, these plasmids had approximately 100% identity and coverage with various *E. coli* plasmids (e.g., pC6-2, pEC1188-MCR, pZE36, pMCR-1_Msc, pDIB-1, pMCR_1139_A1) from strains from various locations around the world, which suggests the horizontal trasmission and origin of these elements. In addition, the *bla*NDM-1 gene was carried in 19 plasmids of 13/24 positive *bla*NDM-1 genomes, mainly in the ST197/Singapore genomes (featuring two plasmids, one conjugative, each carrying a copy of *bla*NDM-1) that also co-harbored the BRP gene (Table S13). Particularly, among the Brazilian genomes (n=20), only 5/11 ST93 genomes (GCA_022170285.1, GCA_022170305.1, GCA_022170345.1, GCA_022170485.1, GCA_022170505.1) had similar plasmids of 74 kb carrying *bla*KPC-2 (Table S13). Interestingly, some integrons (n=21) were present in plasmids, in which CP042532 plasmid harbored the *bla*IMP, *aac, cat*B, *qac* cassette genes; and CP070519 plasmid harbored *qac, arr, cat*B, *aac, bla*OXA-1 cassette genes.

**Figure 3.**
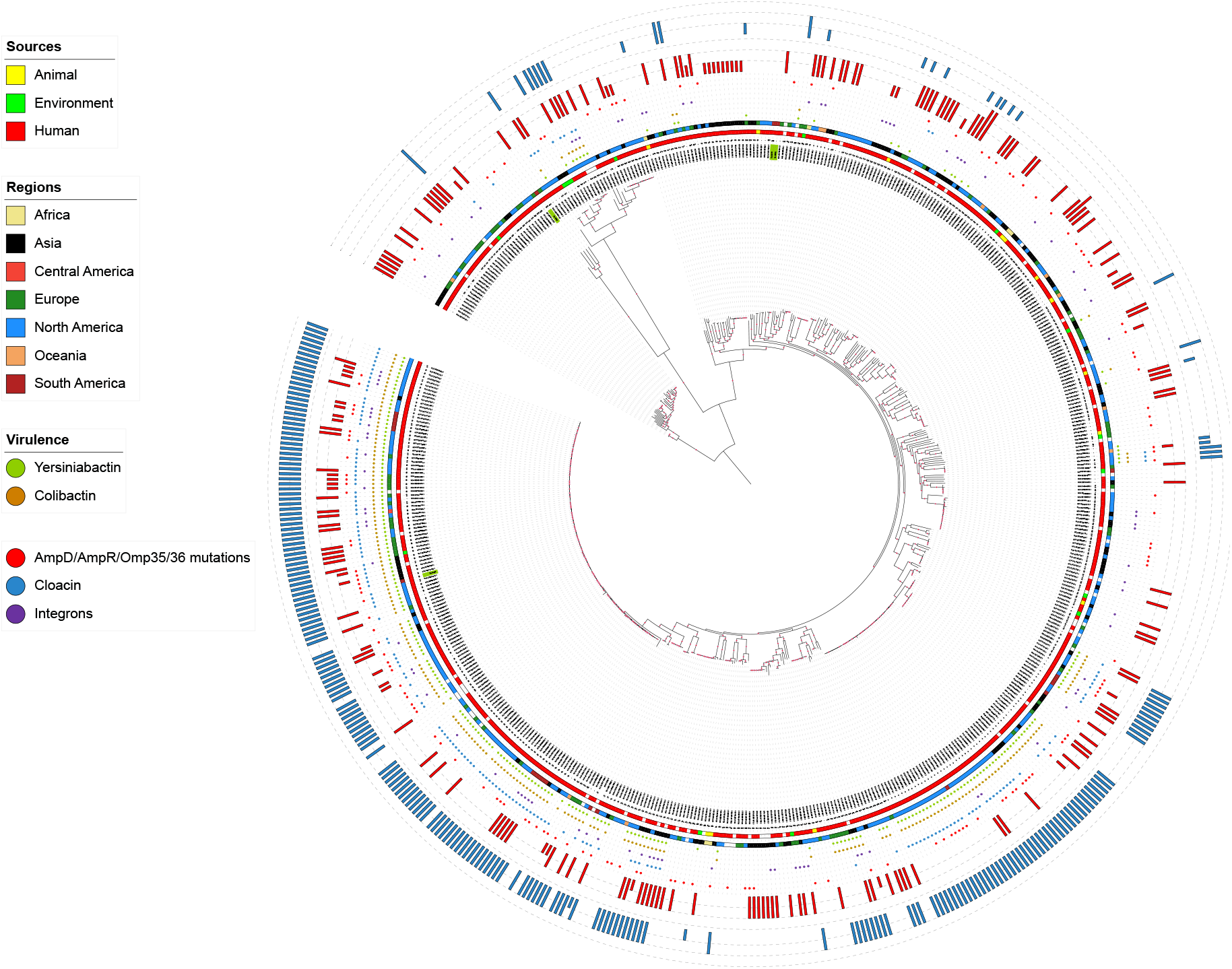
Number and proportion of antibiotic resistance genes in the context of plasmids and chromosomes.

Although most genomes are of human origin, few plasmids from other sources could be observed, some of them carrying resistance genes (e.g., CP050069 from water, carrying *aph, mph*A, *qnr, bla*VIM, *cat, dfr*A, *sul* genes; VKNT01000070 from shower siphon water, carrying *bla*TEM, *aph, bla*KPC genes; CP047668 from chicken manure, carrying *aph, bla*CTX-M-65, *fos*A, *bla*NDM genes).

## Discussion

There is a historical gap in the epidemiology of *K. aerogenes*, as it has always been considered an opportunistic pathogen, but recently interest in this organism has increased, mainly due to the emergence of carbapenem-resistant isolates [3]. Furthermore, this species was recently renamed, being previously known as *Enterobacter aerogenes*, which still influences researchers to keep the old species name, even in recent publications. However, in genomic information databases such as GenBank, this has been updated. Until now, studies focused on this emerging pathogen were small-scale, considering few genomes and/or with a restricted geographic perspective. Here, in addition to generating new genomes of this species in Brazil and contextualizing them in the global scenario, we carried out an in-depth analysis of the resistome, virulome, and mobilome in view of the emergence of *K. aerogenes* as a pathogen.

Previous genomic epidemiological analyzes have shown that *K. aerogenes* ST4 and ST93 have been implicated as the dominant global clones [4], as have occurred in a few countries in the Americas, Europe, and China. In fact, here, expanding this analysis to hundreds of genomes, these STs continue to represent the dominant clones, causing infections/outbreaks in several other countries (e.g., Belgium, Germany, Japan, Qatar, Singapore, South Korea, Switzerland, Thailand, United Kingdom), thus characterizing them as pandemic clones that impac the clinics.

*K. aerogenes* genomes can be clustered by the presence/absence of two virulence determinants: the yersiniabactin and colibactin [5]. In fact, it is observed that most STs lack these loci, while they are predominant in the ST4 and ST93 genomes. These virulence loci were found in several ICEKp, mainly in ICEKp10, and their distribution is not homogeneous among all genomes, even within the same ST, which shows their mobile characteristic. In *K aerogenes*, ICEKp10 had already been identified in a few strains of ST4 and ST93 from the USA [4]. The present analysis revealed a wide distribution of this virulence-associated element within this species, particularly in ST4 and ST93 genomes from several countries around the world. Moreover, colibactin, a genotoxin that can induce DNA damage in eukaryotic cells and tumor formation [25], raises concernings about its high prevalence and distribution in *Klebsiella* species.

In contrast to the *ybt* and *clb* loci, the salmochelin loci (*iro*), another virulence trait identified, was present in a high proportion of genomes (∼92%), including those of animal and environmental origin. However, no genome presented the entire cluster (*iro*BCDEN), lacking *iro*DE. This result is similar to the findings of other studies considering smaller genome sets [5, 26], and suggests that *K. aerogenes* may be a potential reservoir of virulence genes for other bacteria [26]. In fact, in *K. pneumoniae*, salmochelin is detected in low prevalence, being more identified in mobile elements, such as plasmids [27]. In avian pathogenic *Escherichia coli*, it was observed that the absence of *iro*C, *iro*DE, or *iro*N abrogated the virulence of the bacteria [28]. Thus, it can be hypothesized that the *iro* loci in *K. aerogenes* would not act on virulence. However, its widespread presence suggests some role, even if it is different from what occurs in *E. coli*.

Worryingly, a putative virulence island, carrying the aerobactin loci (*iuc*A-D and *iut*A) and *rmp*A2 gene, was found in a *K. aerogenes* genome (GCA_023553355.1). This island has also been observed in several *K. pneumoniae* plasmids. In fact, recently, a transposon harboring the aerobactin operon was identified in *K. pneumoniae* virulence plasmids [29]. Most of these plasmids harbour the *rmp*A/*rmp*A2 genes, aerobactin, and salmochelin loci, which greatly enhance the virulence of *K. pneumoniae* strains [30]. Salmochelin has been observed in almost all *K. aerogenes*, including the GCA_023553355.1 genome. Thus, here we show evidence of horizontal gene transfer of a mobile element carrying hypervirulence marker genes from *K. pneumoniae* to the emergent pathogen *K. aerogenes*.

Although most genomes did not carry acquired antibiotic resistance genes, having only the intrinsic resistome, this does not mean an absence of resistance. Since *in vitro* analyses of Ka-04RR showed its MDR profile, and futher *in silico* analyses revealed the presence of alterations in chromosomally encoded factors associated with this MDR profile. Indeed, chromosomal alterations have been observed in several other genomes. Most genomes (74%; 411/561) do not carry carbapenemase genes (including Ka-04RR), however, almost all of them carry the *amp*C gene, which if overexpressed can lead to carbapenem resistance [3]. In fact, AmpC is chromosomally encoded by *K. aerogenes*, and alterations in its regulatory genes can affect AmpC translation [19]. Here we observed several amino acid replacements in AmpC regulatory proteins, some of them already associated with resistance to carbapenems (mainly in AmpD), in addition to a large set of other replacements yet to be characterized in further studies. Moreover, hundreds of genomes showed truncated Omp35 and Omp36 porins, which can also increase resistance to β-lactam agents [3, 31]. Indeed, a greater number of mutated sites was observed in Omp36 and AmpD proteins, suggesting that their genes are under constant selection. Therefore, regardless of the absence of β-lactamase genes in most genomes, we could infer the prevalence of some level of resistance to β-lactams in *K. aerogenes* circulating around the world.

Unlike *K. pneumoniae*, reports of integrons in *K. aerogenes* are scarce [32-34]. Here, this genetic element was identified in ∼17% of the analyzed genomes, presenting several combinations of gene cassettes, some not yet reported in this species, such as *bla*OXA-1, *bla*OXA10, *bla*IMP, *bla*VIM, *bla*GES. Thus, carbapenemase genes can also be captured by these genetic platforms in this species.

In Brazil, studies on *K. aerogenes* have focused on molecular analyses of resistance genes from carbapenem-resistant isolates. These studies cover isolates from three Brazilian regions (Minas Gerais, Paraná, and Pernambuco states), most of them co-harboring the *bla*KPC-2 and *bla*TEM genes [35-37]. In fact, we observed that several Brazilian of ST4 and ST93 genomes presented these genes (Table S1), and plasmids carrying carbapenemase genes (*bla*KPC-2, *bla*NDM-1) have already been observed among *K. aerogenes* in the country [38, 39]. In addition, we provided new genomes from other regions, expanding regional epidemiology. Although data on resistance genes in *K. aerogenes* in Brazil are available, most studies have not defined the ST of these strains, with only reports of ST93 and ST16 [8, 40]. Thus, the current epidemiological scenario of *K. aerogenes* in Brazil is mainly driven by ST93, ST16 and also by ST4, as is the case worldwide. In addition, a clinical *bla*NDM-producing *K. aerogenes* was recently identified in the country belonging to ST128 [41]. Here, ST128 was also associated with environmental (Asia) and animal (Africa) strains, therefore revealing the One Health trait of this species, particularly of ST128. Furthermore, this ST is widespread and evolving in the context of the acquisition of resistance and virulence genes.

In China, *bla*NDM alleles were associated with plasmids in *K. aerogenes* ST4 [42, 43]. Here, the *bla*NDM gene was also found in several plasmids from Singaporean genomes. Interestingly, both countries were the only ones to present genomes with plasmids carrying the *mcr* gene (colistin resistance). Indeed, the initial report of *mcr* in plasmids occurred in China (2016) [44], and our analysis showed that the presence of the *mcr* gene in plasmids of *K. aerogenes* still seems to be restricted to Asia. Even though these plasmids are already widespread in *E. coli* worldwide. Although plasmids drive the exchange of antibiotic resistance in other countries, we could only observe five Brazilian ST93 genomes harboring plasmids with *bla*KPC-2. Therefore, other transfer mechanisms may be acting in other acquired genes. Nevertheless, if plasmids are also driving *K. aerogenes* adaptation in Brazil, more sequences should be made available to determine this question, in addition to more data for epidemiology.

The most common type of plasmid replicon identified was the ColRNAI, present in hundreds of genomes, as observed by Passarelli-Araujo^a^ et al. (2019) on a smaller set of genomes. Interestingly, hundreds of ColRNAI replicon plasmids have been observed carrying colicin E3 (cloacin-like; rRNase activity) and its immunity gene. Colicins are bacteriocins produced by some gram-negative bacteria, showing antibacterial activity against closely related species, and being highly found in natural populations of *E. coli* [45, 46]. Indeed, blasting cloacin-bearing *K. aerogenes* plasmids, several *E. coli* and *K. pneumoniae* plasmids showed high coverage and identity of >75% and >99%, respectively. It has been reported that Cloacin-like exhibits a weaker effect and narrower spectrum of activity against several species, but exhibits high inhibitory activity against *K. aerogenes*, which suggests that these plasmids are associated with the ecology of this species [46]. Interestingly, most *K. aerogenes* genomes contained the *iut*A gene, which encodes a receptor for aerobactin, a rare virulence factor in the current dataset. Curiously, this receptor is also used by cloacins [46], suggesting that the main function of this receptor in *K. aerogenes* may not be related to virulence, but perhaps to ecological functions.

## Methods

### Public data set

Complete and draft genomes of *Klebsiella aerogenes* (n=557) were obtained from the National Center for Biotechnology Information (NCBI) in October 2022. The accession numbers are supplied in Table S1.

### Genome sequencing and assembly

In this study we generated four *Klebsiella aerogenes* genomes from nosocomial cases in the Amazonic and Southeast regions of Brazil. The genomic DNA extraction was done using the NucleoSpin Microbial DNA kit (Macherey-Nagel), and the genomic libraries were constructed using Nextera paired-end library. The sequencing was performed using Illumina Hiseq 2500, generating reads of 150 bp length. The raw reads were filtered and trimmed using NGS QC Toolkit v.2.3.3 [47] with a Phred score ≥20. The genomes were *de novo* assembled using SPAdes assembler v3.14.1 [48].

### Antimicrobial susceptibility testing

Antimicrobial susceptibility testing was performed by disk-diffusion method using 24 antibiotics (amikacin, gentamicin, streptomycin, tobramycin, neomycin, kanamycin, imipenem, meropenem, ertapenem, ceftazidime, ceftriaxone, cefotaxime, cefepime, cefoxitin, cefuroxime, ampicillin/sulbactam, piperacillin/tazobactam, ticarcillin/clavulanic acid, ciprofloxacin, levofloxacin, trimethoprim/sulfamethoxazole, minocycline, tetracycline, colistin) considered for *Enterobacteriaceae* resistance classification [17] and interpreted according to the Clinical and Laboratory Standards Institute [49]. The minimum inhibitory concentration of colistin was determined by the broth microdilution method and interpreted according to the European Committee for Antimicrobial Susceptibility Testing (EUCAST) guidelines (MIC breakpoint for resistance > 2 mg/L) [50].

### Genome characterization

The *K. aerogenes* genomes were submitted to the Kleborate v2.1.0 [51] pipeline to sequence typing, and identification of acquired virulence and antibiotic resistance genes. This pipeline scores the virulence and antibiotic resistance profiles of the genomes based on the presence of clinically relevant gene markers. The resistance scores range from 0 to 3, based on the presence of genes associated with ESBL (score 1), carbapenemases (score 2), and carbapenemase plus colistin resistance (score 3). The Virulence scores range from 0 to 5, based on the presence of three loci (yersiniabactin, colibactin, and aerobactin): yersiniabactin only (score 1), colibactin without aerobactin (score 2), aerobactin only (score 3), aerobactin and yersiniabactin without colibactin (score 4), and all three present (score 5).

Plasmid identification and characterization was performed with ABRicate v1.0.1 (https://github.com/tseemann/abricate) using PlasmidFinder [52], VFDB [53], and CARD [54] databases. Integrons were survey with Integron Finder [55]. Type VI Secretion System (T6SS) was identified using the T6SS prediction tool [56]. Plasmid clustering was done using CD-HIT-EST at 90% identity and 70% coverage [57]. The mobility of the plasmids was predicted based on the presence of gene markers, such as relaxase, Type IV Secretion System (T4SS) genes (e.g., VirB4 and VirD4), and OriT sequences (origin sites of DNA transfer), as described [24, 58]. The plasmids were classified as: conjugative, if encoding relaxase, *vir*B4, and *vir*D4 homologues; mobilizable, if encoding relaxase and/or oriT sequences, and lacking the T4SS; and non-mobilizable, if not encoding relaxase and/or oriT sequences. These searches were performed using hmm profiles as previous described [58], in addition to the *vir*B4 homologue CagE_TrbE_VirB (Pfam PF03135), present in some Proteobacteria.

### Phylogeny

The genomes were annotated by Prokka v1.12 [59]. *K. aerogenes* core genome was estimated using Roary v3.13 [60], and single nucleotide polymorphisms (SNPs) were extracted from the concatenated core genes with snp-sites v2.5.1 [61]. Next, phylogenetic analysis was performed by IQTree v1.6.12 [62] to obtain a maximum likelihood tree with 1000 ultrafast bootstrap replicates [63]. The tree was designed using iTOL platform [64].

## Acknowledgements

Este estudo foi financiado pela FAPERJ - Fundação Carlos Chagas Filho de Amparo à Pesquisa do Estado do Rio de Janeiro, Processo SEI-260003/019688/2022.

This study was supported by Conselho Nacional de Desenvolvimento Científico e Tecnológico (CNPq) and Oswaldo Cruz Institute Grants.

## Author contributions

Conceptualization, ACV, SM; methodology, SM, ACV; formal analysis, SM; investigation, FF; Resources, RC; writing—original draft preparation, SM and ACV; writing—review and editing, SM, ACV, and EF; supervision, ACV. All authors have read and agreed to the published version of the manuscript.

## Data availability

The data analyzed in this study is available in the supplementary information files. The genomes generated are available under BioProject PRJNA1000993 and GenBank accession numbers: JAUUCT000000000 (Ka-01RR), JAUUCS000000000 (Ka-02RR), JAUUCR000000000 (Ka-04RR), JAUUCQ000000000 (Ka-06RJ).

## Notes

### Competing Interest Statement

The authors have declared no competing interest.

## References

1. Chavda, K. D., Chen, L., Fouts, D. E., Sutton, G., Brinkac, L. et al. Comprehensive Genome Analysis of Carbapenemase-Producing Enterobacter spp.: New Insights into Phylogeny, Population Structure, and Resistance Mechanisms. mBio. 7(6), e02093–16; 10.1128/mBio.02093-16 (2016).

2. Tindall, B. J., Sutton, G., Garrity, G. M. Enterobacter aerogenes Hormaeche and Edwards 1960 (Approved Lists 1980) and Klebsiella mobilis Bascomb et al. 1971 (Approved Lists 1980) share the same nomenclatural type (ATCC 13048) on the Approved Lists and are homotypic synonyms, with consequences for the name Klebsiella mobilis Bascomb et al. 1971 (Approved Lists 1980). Int J Syst Evol Microbiol. 67(2), 502–504; 10.1099/ijsem.0.001572 (2017).

3. Merhi, G., Amayri, S., Bitar, I., Araj, G. F., Tokajian, S. Whole Genome-Based Characterization of Multidrug Resistant Enterobacter and Klebsiella aerogenes Isolates from Lebanon. Microbiol Spectr. 11(1), e0291722; 10.1128/spectrum.02917-22 (2023).

4. Malek, A., McGlynn, K., Taffner, S., Fine, L., Tesini, B. et al. Next-Generation-Sequencing-Based Hospital Outbreak Investigation Yields Insight into Klebsiella aerogenes Population Structure and Determinants of Carbapenem Resistance and Pathogenicity. Antimicrob Agents Chemother. 63(6), e02577–18; 10.1128/AAC.02577-18 (2019).

5. Passarelli-Araujoa, H., Palmeiro, J. K., Moharana, K. C., Pedrosa-Silva, F., Dalla-Costa, L. M. et al. Genomic analysis unveils important aspects of population structure, virulence, and antimicrobial resistance in Klebsiella aerogenes. FEBS J. 286(19), 3797–3810; 10.1111/febs.15005 (2019).

6. Mazumder, R., Hussain, A., Bhadra, B., Phelan, J., Campino, S. et al. Case report: A successfully treated case of community-acquired urinary tract infection due to Klebsiella aerogenes in Bangladesh. Front Med. 10, 1206756; 10.3389/fmed.2023.1206756 (2023).

7. Diene, S. M., Merhej, V., Henry, M., El Filali, A., Roux, V. et al. The rhizome of the multidrug-resistant Enterobacter aerogenes genome reveals how new ‘killer bugs’ are created because of a sympatric lifestyle. Mol Biol Evol. 30(2), 369–383; 10.1093/molbev/mss236 (2013).

8. da Silva, K. E., de Almeida de Souza, G. H., Moura, Q., Rossato, L., Limiere, L. C. et al. Genetic Diversity of Virulent Polymyxin-Resistant Klebsiella aerogenes Isolated from Intensive Care Units. Antibiotics. 11(8), 1127; 10.3390/antibiotics11081127 (2022).

9. Loiwal, V., Kumar, A., Gupta, P., Gomber, S., Ramachandran, V. G. Enterobacter aerogenes outbreak in a neonatal intensive care unit. Pediatr Int. 41(2), 157–161; 10.1046/j.1442-200x.1999.4121033.x (1999).

10. Hao, M., Shen, Z., Ye, M., Hu, F., Xu, X. et al. Outbreak Of Klebsiella pneumoniae Carbapenemase-Producing Klebsiella aerogenes Strains In A Tertiary Hospital In China. Infect Drug Resist. 12, 3283–3290; 10.2147/IDR.S221279 (2019).

11. Narayan, S. A., Kool, J. L., Vakololoma, M., Steer, A. C., Mejia, A. et al. Investigation and control of an outbreak of Enterobacter aerogenes bloodstream infection in a neonatal intensive care unit in Fiji. Infect Control Hosp Epidemiol. 30(8), 797–800; 10.1086/598240 (2009).

12. Baier-Grabner, S., Equiluz-Bruck, S., Endress, D., Blaschitz, M., Schubert, S. et al. A Yersiniabactin-producing Klebsiella aerogenes Strain Causing an Outbreak in an Austrian Neonatal Intensive Care Unit. Pediatr Infect Dis J. 41(7), 593–599; 10.1097/INF.0000000000003553 (2022).

13. Hallbäck, E. T., Johnning, A., Myhrman, S., Studahl, M., Hentz, E. et al. Outbreak of OXA-48-producing Enterobacteriaceae in a neonatal intensive care unit in Western Sweden. Eur J Clin Microbiol Infect Dis. 42(5), 597–605; 10.1007/s10096-023-04584-y (2023).

14. Piagnerelli, M., Kennes, B., Brogniez, Y., Deplano, A., Govaerts, D. Outbreak of nosocomial multidrug-resistant Enterobacter aerogenes in a geriatric unit: failure of isolation contact, analysis of risk factors, and use of pulsed-field gel electrophoresis. Infect Control Hosp Epidemiol. 21(10), 651–653; 10.1086/501704 (2000).

15. Neuwirth, C., Siebor, E., Lopez, J., Pechinot, A., Kazmierczak, A. Outbreak of TEM-24-producing Enterobacter aerogenes in an intensive care unit and dissemination of the extended-spectrum beta-lactamase to other members of the family enterobacteriaceae. J Clin Microbiol. 34(1), 76–79; 10.1128/jcm.34.1.76-79.1996 (1996).

16. Cheng, Q., Ma, Z., Gong, Z., Liang, Y., Guo, J. et al. Whole-genome sequencing analysis of Klebsiella aerogenes among men who have sex with men in Guangzhou, China. Front Microbiol. 14, 1102907; 10.3389/fmicb.2023.1102907 (2023).

17. Magiorakos, A. P., Srinivasan, A., Carey, R. B., Carmeli, Y., Falagas, M. E et al. Multidrug-resistant, extensively drug-resistant and pandrug-resistant bacteria: an international expert proposal for interim standard definitions for acquired resistance. Clin Microbiol Infect. 18(3):268–281; 10.1111/j.1469-0691.2011.03570.x (2012).

18. Bedenić, B., Vranić-Ladavac, M., Venditti, C., Tambić-Andrašević, A. Barišić, et al. Emergence of colistin resistance in Enterobacter aerogenes from Croatia. Chemother. 30(2), 120–123; doi.org/10.1080/1120009X.2017.1382121 (2018).

19. D’Souza, R., Nguyen, L. P., Pinto, N. A., Lee, H., Vu, T. N. et al. Role of AmpG in the resistance to β-lactam agents, including cephalosporins and carbapenems: candidate for a novel antimicrobial target. Ann Clin Microbiol Antimicrob. 20(1), 45; 10.1186/s12941-021-00446-7 (2021).

20. Kuga, A., Okamoto, R., Inoue, M. ampR gene mutations that greatly increase class C beta-lactamase activity in Enterobacter cloacae. Antimicrob Agents Chemother. 44(3), 561–567; 10.1128/AAC.44.3.561-567.2000 (2000).

21. Caille, O., Zincke, D., Merighi, M., Balasubramanian, D., Kumari, H. et al. Structural and functional characterization of Pseudomonas aeruginosa global regulator AmpR. J Bacteriol. 196(22), 3890–3902; 10.1128/JB.01997-14 (2014).

22. Dupont, H., Choinier, P., Roche, D., Adiba, S., Sookdeb, M. et al. Structural Alteration of OmpR as a Source of Ertapenem Resistance in a CTX-M-15-Producing Escherichia coli O25b:H4 Sequence Type 131 Clinical Isolate. Antimicrob Agents Chemother. 61(5), e00014–17; 10.1128/AAC.00014-17 (2017).

23. Ching, C., Zaman, M. H. Identification of Multiple Low-Level Resistance Determinants and Coselection of Motility Impairment upon Sub-MIC Ceftriaxone Exposure in Escherichia coli. mSphere. 6(6), e0077821; 10.1128/mSphere.00778-21 (2021).

24. Smillie, C., Garcillán-Barcia, M. P., Francia, M. V., Rocha, E. P., de la Cruz F. Mobility of plasmids. Microbiol Mol Biol Rev. 74(3), 434–452; 10.1128/MMBR.00020-10 (2010).

25. Wernke, K. M., Xue, M., Tirla, A., Kim, C. S., Crawford, J. M. et al. Structure and bioactivity of colibactin. Bioorg Med Chem Lett. 30(15), 127280; 10.1016/j.bmcl.2020.127280 (2020).

26. Spadar, A., Perdigão, J., Campino, S., Clark, T. G. Large-scale genomic analysis of global Klebsiella pneumoniae plasmids reveals multiple simultaneous clusters of carbapenem-resistant hypervirulent strains. Genome Med. 15(1), 3; 10.1186/s13073-023-01153-y (2023).

27. Lam, M. M. C., Wyres, K. L., Judd, L. M., Wick, R. R., Jenney, A. et al. Tracking key virulence loci encoding aerobactin and salmochelin siderophore synthesis in Klebsiella pneumoniae. Genome Med. 10(1), 77; 10.1186/s13073-018-0587-5 (2018).

28. Caza, M., Lépine, F., Milot, S., Dozois, C. M. Specific roles of the iroBCDEN genes in virulence of an avian pathogenic Escherichia coli O78 strain and in production of salmochelins. Infect Immun. 76(8), 3539–3549; 10.1128/IAI.00455-08 (2008).

29. Kaspersen, H., Franklin-Alming, F. V., Hetland, M. A. K., Bernhoff, E., Löhr, I. H. et al. Highly conserved composite transposon harbouring aerobactin iuc3 in Klebsiella pneumoniae from pigs. Microb Genom. 9(2), mgen000960; 10.1099/mgen.0.000960 (2023).

30. Fan, L. P., Yu, Y., Huang, S., Liao, W., Huang, Q. S. et al. Genetic characterization and passage instability of a novel hybrid virulence plasmid in a ST23 hypervirulent Klebsiella pneumoniae. Front Cell Infect Microbiol. 12, 870779; 10.3389/fcimb.2022.870779 (2022).

31. Davin-Regli, A., Pagès, J. M. Enterobacter aerogenes and Enterobacter cloacae; versatile bacterial pathogens confronting antibiotic treatment. Front Microbiol. 6, 392; 10.3389/fmicb.2015.00392 (2015).

32. Ploy, M. C., Courvalin, P., Lambert, T. Characterization of In40 of Enterobacter aerogenes BM2688, a class 1 integron with two new gene cassettes, cmlA2 and qacF. Antimicrob Agents Chemother. 42(10), 2557–2563; 10.1128/AAC.42.10.2557 (1998).

33. Wu, K., Wang, F., Sun, J., Wang, Q., Chen, Q. et al. Class 1 integron gene cassettes in multidrug-resistant Gram-negative bacteria in southern China. Int J Antimicrob Agents. 40(3), 264–267; 10.1016/j.ijantimicag.2012.05.017 (2012).

34. Cheng, C., Sun, J., Zheng, F., Lu, W., Yang, Q. et al. New structures simultaneously harboring class 1 integron and ISCR1-linked resistance genes in multidrug-resistant Gram-negative bacteria. BMC Microbiol. 16, 71; 10.1186/s12866-016-0683-x (2016).

35. Tuon, F. F., Scharf, C., Rocha, J. L., Cieslinsk, J., Becker, G. N. et al. KPC-producing Enterobacter aerogenes infection. Braz J Infect Dis. 19(3), 324–327; 10.1016/j.bjid.2015.01.003 (2015).

36. Pereira, R. S., Dias, V. C., Ferreira-Machado, A. B., Resende, J. A., Bastos, A. N. et al. Physiological and molecular characteristics of carbapenem resistance in Klebsiella pneumoniae and Enterobacter aerogenes. Infect Dev Ctries. 10(6), 592–599; 10.3855/jidc.6821 (2016).

37. Cabral, A. B., Maciel, M. A. V., Barros, J. F., Antunes, M. M., Barbosa de Castro, C. M. M. et al. Clonal spread and accumulation of β-lactam resistance determinants in Enterobacter aerogenes and Enterobacter cloacae complex isolates from infection and colonization in patients at a public hospital in Recife, Pernambuco, Brazil. J Med Microbiol. 66(1), 70–77; 10.1099/jmm.0.000398 (2017).

38. Bispo Beltrão, E. M., de Oliveira, É. M., Dos Santos Vasconcelos, C. R., Cabral, A. B., Rezende, A. M. et al. Multidrug-resistant Klebsiella aerogenes clinical isolates from Brazil carrying IncQ1 plasmids containing the blaKPC-2 gene associated with non-Tn4401 elements (NTEKPC-IId). J Glob Antimicrob Resist. 22, 43–44; 10.1016/j.jgar.2020.05.001 (2020).

39. Soares, C. R. P., Oliveira-Júnior, J. B., Firmo, E. F. First report of a blaNDM-resistant gene in a Klebsiella aerogenes clinical isolate from Brazil. Rev Soc Bras Med Trop. 54, e02622020; 10.1590/0037-8682-0262-2020 (2020).

40. Passarelli-Araujob, H., Palmeiro, J. K., Moharana, K. C., Pedrosa-Silva, F., Dalla-Costa, L. M. et al. Molecular epidemiology of 16S rRNA methyltransferase in Brazil: RmtG in Klebsiella aerogenes ST93 (CC4). An Acad Bras Cienc. 91(suppl 1), e20180762; 10.1590/0001-376520182018762 (2019).

41. Camargo, C. H., Yamada, A. Y., Souza, A. R., Reis, A. D., Santos, M. B. N. et al. Genomic Diversity of NDM-Producing Klebsiella Species from Brazil, 2013-2022. Antibiotics (Basel). 11(10), 1395; 10.3390/antibiotics11101395 (2022).

42. Pan, F., Xu, Q., Zhang, H. Emergence of NDM-5 Producing Carbapenem-Resistant Klebsiella aerogenes in a Pediatric Hospital in Shanghai, China. Front Public Health. 9, 621527; 10.3389/fpubh.2021.621527 (2021).

43. Li, Y., Wang, Q., Xiao, X., Li, R., Wang, Z. Emergence of blaNDM-9-bearing tigecycline-resistant Klebsiella aerogenes of chicken origin. J Glob Antimicrob Resist. 26, 66–68; 10.1016/j.jgar.2021.04.028 (2021).

44. Liu, Y. Y., Wang, Y., Walsh, T. R., Yi, L. X., Zhang, R., Spencer, J. et al. Emergence of plasmid-mediated colistin resistance mechanism MCR-1 in animals and human beings in China: a microbiological and molecular biological study. Lancet Infect Dis. 16(2), 161–168; 10.1016/S1473-3099(15)00424-7 (2016).

45. Simons, A., Alhanout, K., Duval, R. E. Bacteriocins, Antimicrobial Peptides from Bacterial Origin: Overview of Their Biology and Their Impact against Multidrug-Resistant Bacteria. Microorganisms. 8(5), 639; 10.3390/microorganisms8050639 (2020).

46. Le, M. N., Nguyen, T. H., Trinh, V. M., Nguyen, T. P., Kawada-Matsuo, M. et al. Comprehensive Analysis of Bacteriocins Produced by the Hypermucoviscous Klebsiella pneumoniae Species Complex. Microbiol Spectr. 11(3), e0086323; 10.1128/spectrum.00863-23 (2023).

47. Patel, R. K., Jain, M. NGS QC Toolkit: A toolkit for quality control of next generation sequencing data. PLoS ONE. 7:e30619; 10.1371/journal.pone.0030619 (2012).

48. Bankevich, A., Nurk, S., Antipov, D., Gurevich, A. A., Dvorkin, M. et al. SPAdes: A new genome assembly algorithm and its applications to single-cell sequencing. J. Comput. Biol. 19:455–477; 10.1089/cmb.2012.0021 (2012).

49. CLSI. Performance Standards for Antimicrobial Susceptibility Testing—Thirty-First Edition: M100. 2021.

50. EUCAST. Breakpoint tables for interpretation of MICs and zone diameters (Version 11.0. 2021). https://www.eucast.org/fileadmin/src/media/PDFs/EUCAST_files/Breakpoint_tables/v_11.0_Breakpoint_Tables.pdf [assessed 13 December 2022].

51. Lam, M. M. C., Wick, R. R., Watts, S. C., Cerdeira, L. T. & Wyres, K. L et al. A genomic surveillance framework and genotyping tool for Klebsiella pneumoniae and its related species complex. Nat Commun. 12(1), 4188; 10.1038/s41467-021-24448-3 (2021).

52. Carattoli, A., Zankari, E., García-Fernández, A., Voldby Larsen, M., Lund, O. et al. In silico detection and typing of plasmids using PlasmidFinder and plasmid multilocus sequence typing. Antimicrob Agents Chemother. 58(7), 3895–3903; 10.1128/AAC.02412-14 (2014).

53. Chen, L., Yang, J., Yu, J., Yao, Z., Sun, L., et al. VFDB: a reference database for bacterial virulence factors. Nucleic Acids Res. 33, D325–D328; 10.1093/nar/gki008 (2005).

54. Alcock, B. P., Raphenya, A. R., Lau, T. T. Y., Tsang, K. K., Bouchard, M., et al. CARD 2020: antibiotic resistome surveillance with the comprehensive antibiotic resistance database. Nucleic Acids Res. 48(D1), D517–D525; 10.1093/nar/gkz935 (2020).

55. Néron, B., Littner, E., Haudiquet, M., Perrin, A., Cury, J., et al. IntegronFinder 2.0: Identification and Analysis of Integrons across Bacteria, with a Focus on Antibiotic Resistance in Klebsiella. Microorganisms. 10(4), 700; 10.3390/microorganisms10040700 (2022)

56. Zhang, J., Guan, J., Wang, M., Li, G., Djordjevic, M., et al. SecReT6 update: a comprehensive resource of bacterial Type VI Secretion Systems. Sci China Life Sci. 66(3), 626–634; 10.1007/s11427-022-2172-x (2023).

57. Li, W., & Godzik, A. Cd-hit: a fast program for clustering and comparing large sets of protein or nucleotide sequences. Bioinformatics. 22(13), 1658–1659; 10.1093/bioinformatics/btl158 (2006).

58. Morgado, S. M., & Vicente, A. C. P. Comprehensive in silico survey of the Mycolicibacterium mobilome reveals an as yet underexplored diversity. Microb Genom. 7(3), mgen000533; 10.1099/mgen.0.000533 (2021).

59. Seemann T. Prokka: Rapid prokaryotic genome annotation. Bioinformatics. 30:2068–2069; 10.1093/bioinformatics/btu153 (2014).

60. Page, A. J., Cummins, C. A., Hunt, M., Wong, V. K., Reuter, S. et al. Roary: Rapid large-scale prokaryote pan genome analysis. Bioinformatics. 31:3691–3693; 10.1093/bioinformatics/btv421 (2015).

61. Page, A. J., Taylor, B., Delaney, A. J., Soares, J., Seemann, T. et al. SNP-sites: Rapid efficient extraction of SNPs from multi-FASTA alignments. Microb. Genom. 2:e000056; 10.1099/mgen.0.000056 (2016).

62. Nguyen, L. T., Schmidt, H. A., von Haeseler, A., Minh, B. Q. IQ-TREE: A fast and effective stochastic algorithm for estimating maximum-likelihood phylogenies. Mol. Biol. Evol. 32:268–274; 10.1093/molbev/msu300 (2015).

63. Minh, B. Q., Nguyen, M. A., von Haeseler, A. Ultrafast approximation for phylogenetic bootstrap. Mol. Biol. Evol. 30:1188–1195; 10.1093/molbev/mst024 (2013).

64. Letunic, I., Bork, P. Interactive Tree Of Life (iTOL) v4: Recent updates and new developments. Nucleic Acids Res. 47:W256–W259; 10.1093/nar/gkz239 (2019).

